# Primary neurons can enter M-phase

**DOI:** 10.1101/288589

**Authors:** Chaska C Walton, Wei Zhang, Iris Patiño-Parrado, Estíbaliz Barrio-Alonso, Juan-José Garrido, José M Frade

## Abstract

Differentiated neurons can undergo cell cycle re-entry during pathological conditions, but it remains largely accepted that M-phase is prohibited in these cells. Here we show that primary neurons at post-synaptogenesis stages of development can enter M-phase. We induced cell cycle re-entry by overexpressing a truncated Cyclin E isoform fused to Cdk2. Cyclin E/Cdk2 expression elicits canonical cell cycle checkpoints, which arrest cell cycle progression and trigger apoptosis. As in mitotic cells, checkpoint abrogation enables cell cycle progression through S and G2-phases into M-phase. Although most neurons enter M-phase, only a small subset undergo cell division. Alternatively, neurons can exit M-phase without cell division and recover the axon initial segment, a structural determinant of neuronal viability. We conclude that neurons and mitotic cells share S, G2 and M-phase regulation.

---

Neurons that are not fully differentiated can enter M-phase and undergo cell division^1–10^. In contrast, fully differentiated neurons do not undergo M-phase entry and cell division upon acute induction of cell cycle re-entry^1,11–14^. In pathologies such as Alzheimer’s disease (AD), Parkinson’s disease (PD), amyotrophic lateral sclerosis (AML) or brain injury, neuronal cell cycle re-entry is associated to increased susceptibility to cell death instead of cell division^15,16^. This observation has led to suggest that M-phase entry is prohibited in neurons^16^, and that the cell cycle machinery becomes pro-apoptotic in these cells^17^. However, the neuron-specific mechanisms that block M-phase entry remain unidentified. Furthermore, whether M-phase entry is irreversibly prohibited remains to be determined as well.

The block on M-phase entry could be explained by the presence of canonical cell cycle checkpoints. In mitotic cells, non-physiological cell cycle re-entry activates checkpoints that arrest the cell cycle^18–20^ and can result in cell death to prevent potentially cancerous cell division^18,21^. Cell cycle checkpoint abrogation in mitotic cells can prevent cell death, and enable M-phase entry and cell division^18,19,22^. This suggests that, by abrogating cell cycle checkpoint activity, neuronal M-phase entry and cell division in neurons that undergo cell cycle re-entry should be possible. This possibility remains untested.

To study whether cell cycle checkpoints regulate cell cycle progression in neurons as in mitotic cells, we induced neuronal cell cycle re-entry with a low molecular weight (LMW) Cyclin E isoform (Cyclin ET1)^23^ fused to Cdk2 (t1EK2). t1EK2 overexpression was coupled with genetic and pharmacological checkpoint signaling abrogation. We assessed cell cycle progression through each of its phases. We show that the regulation of S, G2 and M phases in neurons is as in standard mitotic cells. Neurons readily enter M-phase and a small subset can undergo cell division. We also assessed the integrity of the axon initial segment (AIS) after M-phase exit without cell division in multinucleated neurons. We show that multinucleated neurons recover the AIS, indicating that aberrant cell cycle re-entry is not necessarily fatal.

## Results

### t1EK2 induces DNA synthesis in differentiated neurons

Cyclin E is the canonical late G1 cyclin that triggers transition into S-phase by activating Cyclin-dependent kinase 2 (Cdk2)^24^ and is necessary for cell cycle re-entry from quiescence^25^. Strikingly, Cyclin E is highly expressed in neurons under physiological conditions^26^, and Cyclin E upregulation is associated to aberrant neuronal cell cycle re-entry^14,27–33^ and in AD^34,35^. Under physiological conditions, Cyclin E forms catalytically inactive complexes with Cdk5 to promote synapse maturation^26^. However, Cdk5 deregulation is associated to neuron diseases^36^. To avoid interfering with endogenous Cdk5 signaling by off target binding of Cyclin ET1 to Cdk5, we generated a t1EK2 fusion product and used it to induce neuronal cell cycle re-entry.

t1EK2 or control LacZ were co-lipofected with red fluorescent protein (RFP) in hippocampal cultures maintained for 15 days *in vitro* (DIV), a stage in which synapses have already been developed^37^. Transfected neurons were identified by MAP2-specific labeling in RFP-positive cells. We studied cell cycle S-phase entry by assessing 5-bromo-2´-deoxyuridine (BrdU) incorporation 1, 1.5, and 2 days post-transfection (dpt). Transfected control neurons never incorporated BrdU (n=491) (Fig. 1a, b), confirming that neurons in primary culture at these stages of maturation do not spontaneously re-enter the cell cycle. In contrast, t1EK2 induced DNA synthesis in neurons (Fig. 1a, b). At 1 dpt, 52.3 *%* t1EK2-expressing neurons incorporated BrdU (*t*_152_=12.906, p<0.001). This increased at 1.5 dpt to 58.9 *%* (*t*_128_=13.548, p<0.001), and then declined to 46.8 *%* at 2 dpt (*t*_93_=9.047, p<0.001). The apparent decrease in BrdU incorporation from 1.5 to 2 dpt likely reflected selective death of t1EK2 transfected neurons since, at 3 dpt, BrdU incorporation could no longer be assessed due to widespread deterioration of the t1EK2-expressing neurons.

**Fig. 1.**
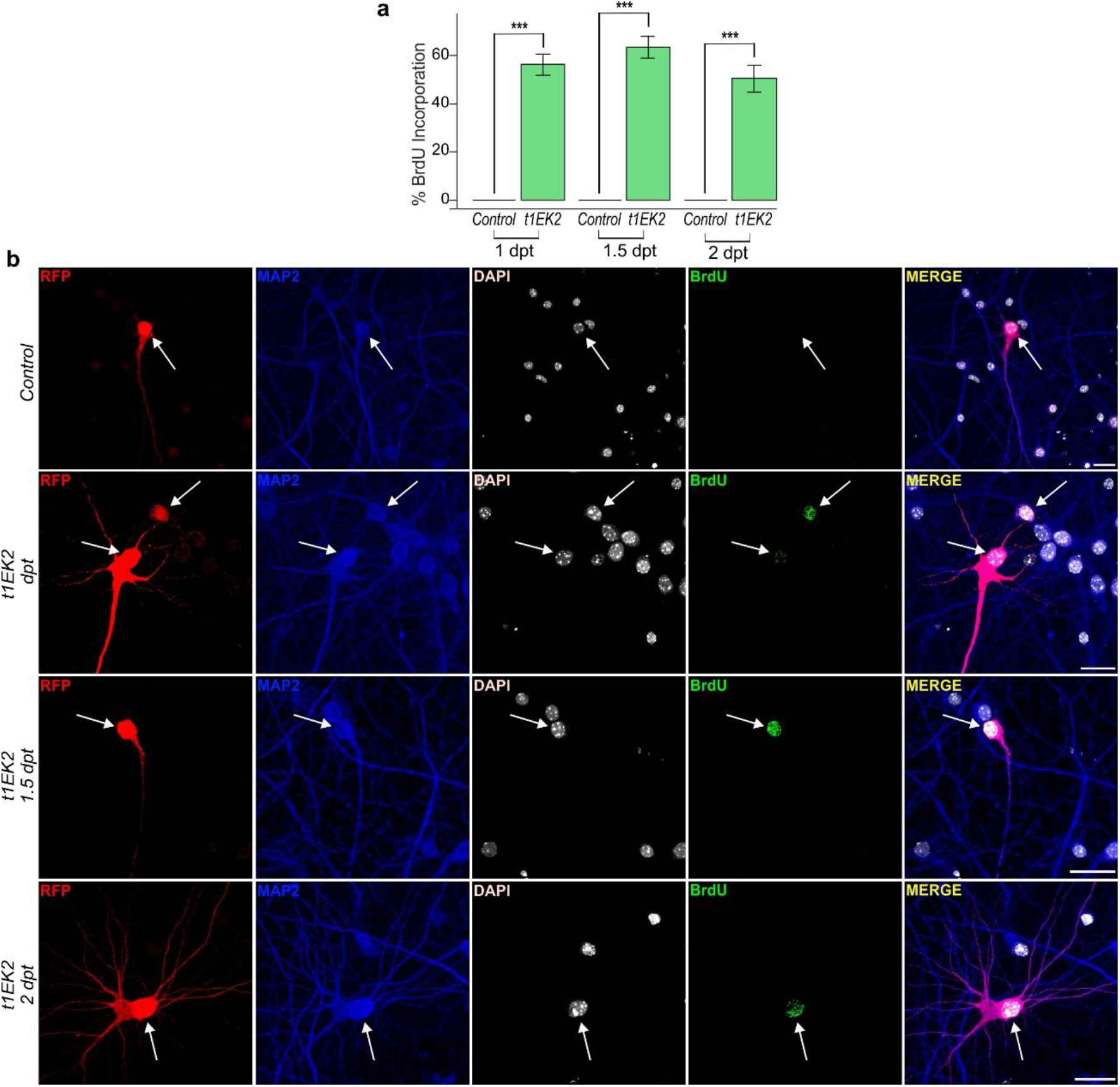
t1EK2 induces DNA synthesis in differentiated neurons. **a** Percentage of BrdU positive neurons expressing LacZ (Control) or t1EK2 at 1 (Control: n=129; t1EK2: n=153), 1.5 (Control: n=201; t1EK2: n=129), and 2 (Control: n=161; t1EK2: n=94) dpt (16-17 DIV). Control neurons never incorporate BrdU. Three independent experiments for each dpt were carried out. As control neurons never incorporated BrdU, statistical analysis was performed by comparing percent BrdU incorporation in t1EK2-expressing neurons with zero (***p<0.001; one-tailed *t* test for each dpt). BrdU incorporation could not be assessed at 3 dpt due to widespread neuronal death of t1EK2-expressing neurons. **b** Representative confocal images of BrdU incorporation in a LacZ control neuron (1 dpt) and t1EK2-expressing neurons at 1, 1.5 and 2 dpt. Graphs represent mean ± s.e.m. Scale bar: 25 μm.

### t1EK2 induces p53-dependent apoptosis in differentiated neurons

Cell cycle related cell death was expected, as Cyclin E is consistently associated to apoptotic cell death during aberrant neuronal cell cycle re-entry^14,27,29,31–33^. To confirm whether t1EK2 was inducing apoptosis we performed active caspase-3 immunolabeling^38^. The proportion of neurons expressing t1EK2 that were positive for active caspase-3 (34.7 %) was already significantly higher than control neurons (6.0 %) at 1.5 dpt (p<0.001, Fisher’s exact test) (Fig. 2a). As expected, apoptotic neurons displayed pyknotic nuclei (Fig. 2b). We therefore concluded that t1EK2 induced neuronal S-phase entry and this was followed by apoptotic cell death.

**Fig. 2.**
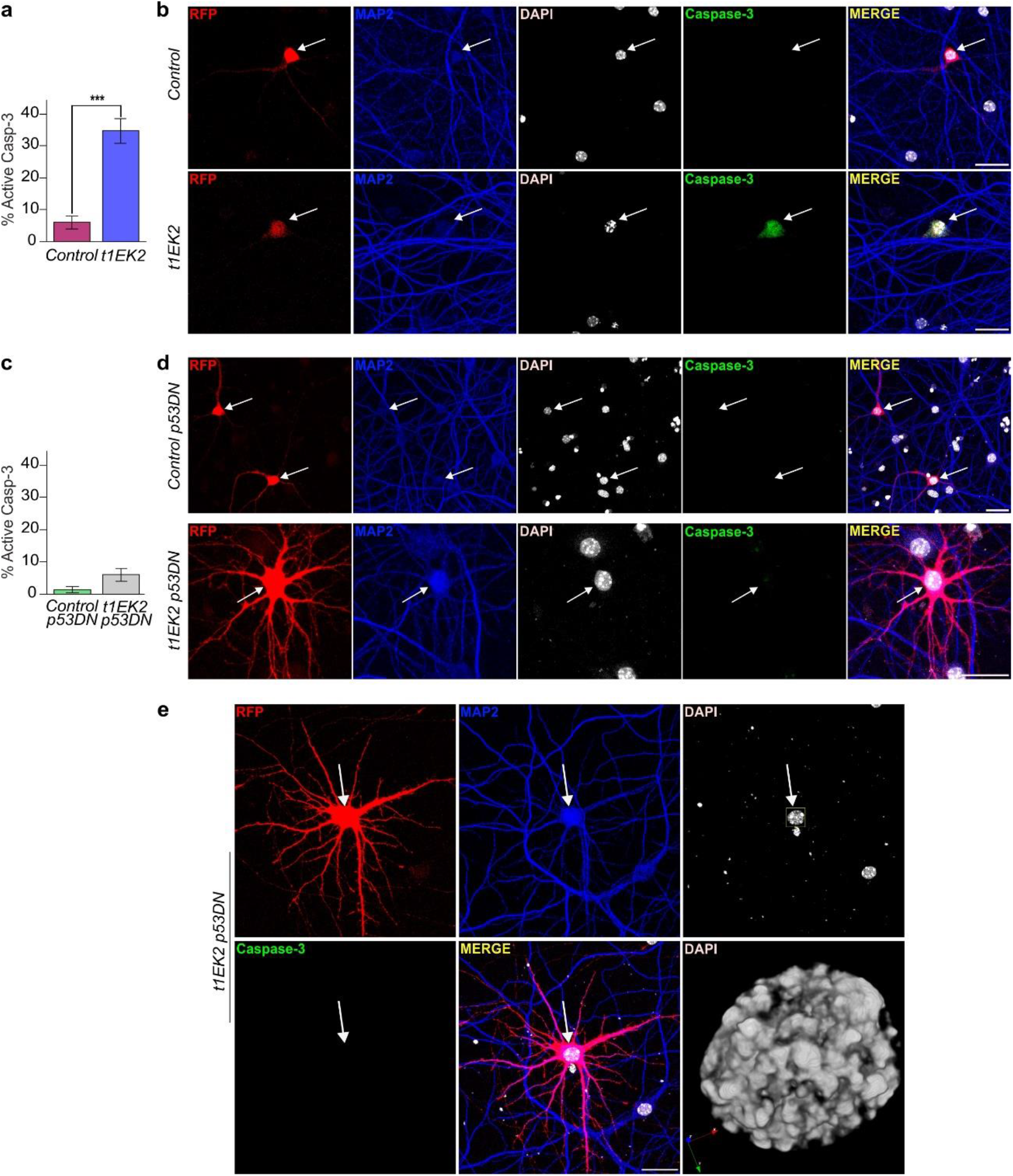
t1EK2 induces p53-dependent apoptosis in differentiated neurons. **a** Percentage of active caspase-3 positive neurons expressing LacZ (n=150) or t1EK2 (n=150) at 1.5 dpt (17 DIV). Three independent experiments were carried out (***p<0.001; two-sided Fisher’s exact test). **b** Representative confocal images of active caspase-3-specific immunostaining in a LacZ control neuron and a t1EK2-expressing neuron. Note that the active caspase-3-positive nuclei is pyknotic. Arrows identify RFP positive neurons. **c** Percentage of active caspase-3 positive neurons expressing LacZ/p53DN (n=150) or t1EK2/p53DN (n=150) at 1.5 dpt (17 DIV). Three independent experiments were carried out (***p<0.001; two-sided Fisher’s exact test). **d** Representative confocal images of active caspase-3-negative LacZ/p53DN (Control) and t1EK2/p53DN expressing neurons. **e** Confocal image of an active caspase-3-negative neuron presenting mitotic chromatin condensation characteristic of prophase with 3D reconstruction of the nucleus. Blue, red and green arrows represent 3D orientation. Graphs represent mean ± s.e.m. Scale bar: 25 μm.

A key barrier to Cyclin E/Cdk2 deregulation is the p53 tumor suppressor protein^39^. It is therefore possible that p53 activity could underlie the apoptotic response of hippocampal neurons to t1EK2-induced cell cycle re-entry. In consequence, cell cycle-related cell death in t1EK2-expressing neurons should be blocked by suppressing p53 signaling. To test this hypothesis, we prevented p53 function in t1EK2-expressing neurons by expressing a dominant negative mutant (p53DN)^40^. We assessed apoptosis using active caspase-3 immunostaing. p53 blockade prevented cell cycle re-entry-induced apoptosis in neurons (Fig. 2c-e). The proportion of active caspase-3-positive t1EK2/p53DN-expressing neurons (6.0 %) was not statistically significantly different from control neurons expressing p53DN (1.3 %) at 1.5 dpt (p=0.061, Fisher’s exact test) (Fig. 2c). Moreover, rescued neurons occasionally showed mitotic chromatin condensation (Fig. 2e), evidencing that t1EK2 overexpression plus p53 blockade can enable neuronal cell cycle progression into M-phase.

### p53 tumor suppressor blockade rescues t1EK2-induced BrdU incorporation

We quantified BrdU incorporation at 1, 2, and 3 dpt to confirm that preventing p53-dependent apoptosis rescued BrdU-positive neurons. As we could not analyze BrdU incorporation at 3 dpt when expressing t1EK2 without p53DN (Fig. 1a), we included this time point to determine whether the effects of p53DN were sustained. Control neurons never incorporated BrdU (n=1,043) (Fig. 3a, b). In contrast, a substantial proportion of t1EK2/p53DN-expressing neurons was positive for BrdU labelling at 1 (38.8 %) (*t*_375_=15.429, p<0.001), 2 (61.2 %) (*t*_293_=21.509, p<0.001), and 3 dpt (61.8 %) (*t*_203_=18.109, p<0.001) (Fig. 3a, b). Co-expressing p53DN with t1EK2 rescued BrdU incorporation at 3 dpt and, although we did not quantify it, neurons with BrdU incorporation could still be found at 5 dpt (Fig. 3c). In summary, p53 tumor suppressor loss of function prevented t1EK2-induced apoptosis and rescued S-phase viability.

**Fig. 3.**
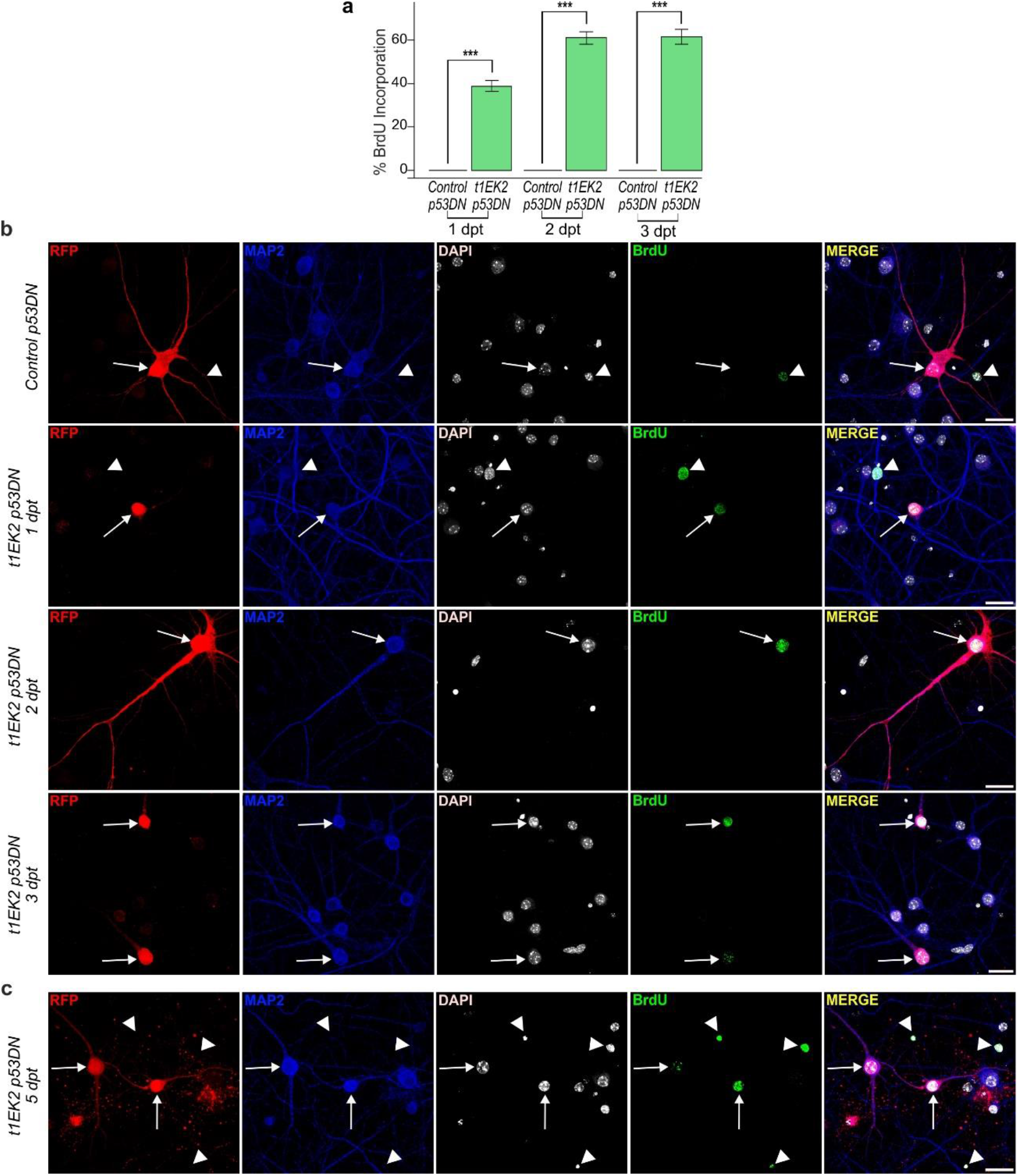
p53 tumor suppressor blockade rescues t1EK2-induced BrdU incorporation. **a** Percentage of BrdU positive neurons expressing LacZ/p53DN (Control) or t1EK2/p53DN at 1 (Control: n=325; t1EK2/p53DN: n=376), 2 (Control: n=406; t1EK2/p53DN: n=294), and 3 (Control: n=312; t1EK2/p53DN: n=204) dpt (16-18 DIV). Control neurons never incorporate BrdU. Three independent experiments for each dpt were carried out. As control neurons never incorporated BrdU, statistical analysis was performed by comparing percent BrdU incorporation in t1EK2/p53DN-expressing neurons with zero (***p<0.001; one-tailed *t* test for each dpt). **b** Representative confocal images of BrdU incorporation of a LacZ/p53DN control neuron (3 dpt) and t1EK2/p53DN-expressing neurons at 1, 2, and 3 dpt. **c** Confocal images showing BrdU incorporation in t1EK2/p53DN-expressing neurons at 5 dpt. Arrows identify RFP positive neurons and arrowheads MAP2-negative (non-neuronal) cells. Graphs represent mean ± s.e.m. Scale bar: 25 μm.

### Wee1 inhibition enables G2/M transition in differentiated neurons

Next, we wanted to determine whether neurons could continue cell cycle progression beyond S-phase. To assess this possibility, we performed Phospho-Ser10 Histone H3 (pH3) immunostaining to identify t1EK2-expressing neurons in late G2 and M-phase^41^. The onset of pH3 staining starts at late G2, prior to M-phase, in the form of nuclear pH3 foci. This transitions to pan-nuclear pH3 staining in M-phase and remains from prophase to metaphase. However, the loss of pH3 staining begins while cells are still in M-phase, at anaphase onset. Thus, we blocked anaphase onset with the proteasome inhibitor MG132^42^ to identify all neurons that entered M-phase (see Fig. S1a, b for protocol details). Consistent with the absence of spontaneous DNA replication, control MAP2-positive neurons were never positive for pH3 (n=150) (Fig. 4a). In contrast, 45.3 % (*t*_149_=11.116, p<0.001) of t1EK2/p53DN-expressing neurons reached late G2 (evidenced by pH3-specific foci). In addition, 2.7 % (*t*_149_=2.020, p<0.05) of t1EK2/p53DN-expressing neurons progressed beyond G2 into M-phase (characterized by pan-nuclear pH3 labelling) (Fig. 4a), an observation consistent with the capacity of active caspase-3-negative neurons expressing t1EK2/p53DN to occasionally enter M-phase (Fig. 2e). Therefore, although most neurons could not undergo G2/M transition, some neurons did enter M-phase upon t1EK2 expression and p53 inhibition.

**Fig. 4.**
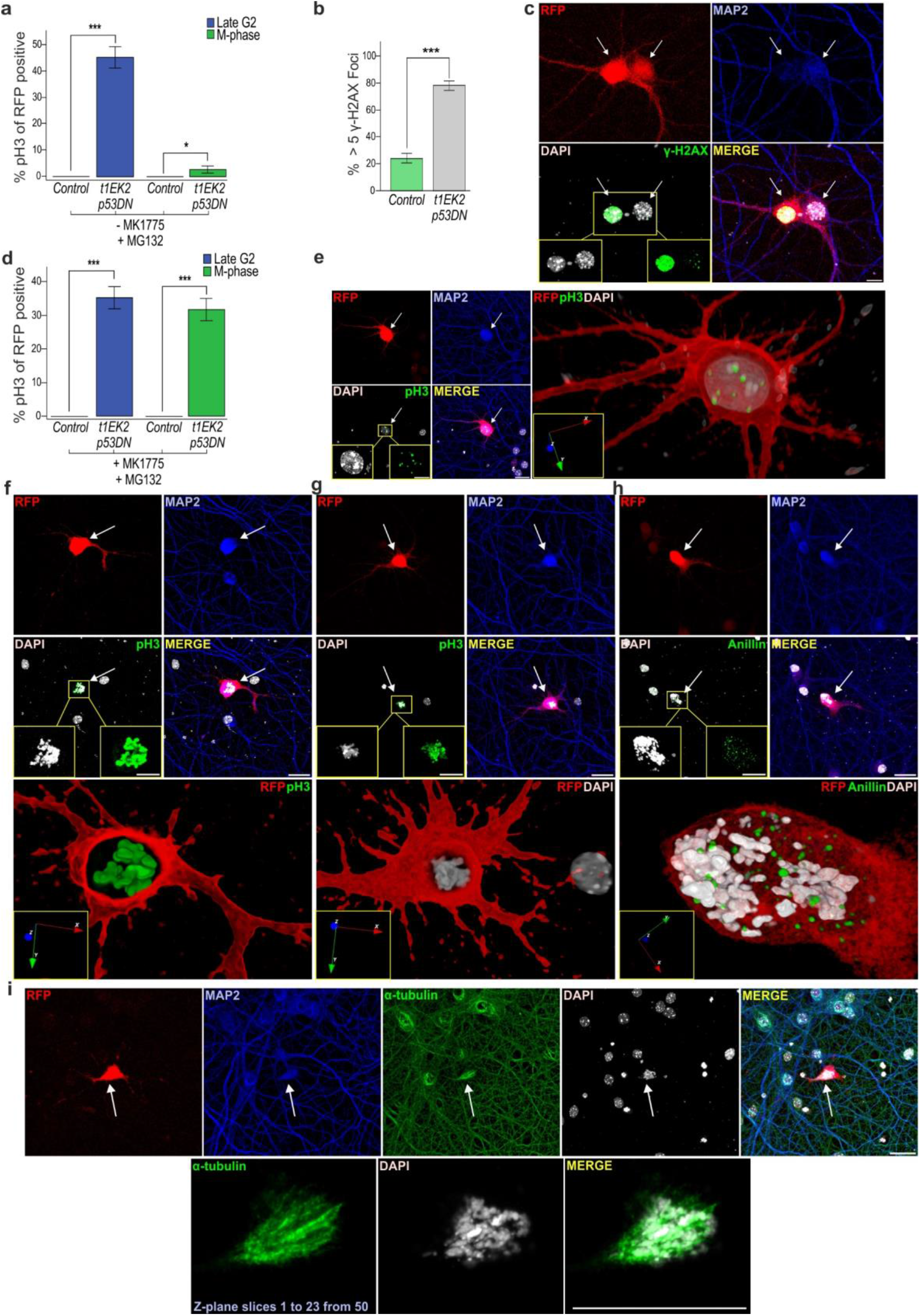
Wee1 inhibition enables G2/M transition in differentiated neurons. **a** Percentage of LacZ/p53DN- (Control) or t1EK2/p53DN-expressing neurons with pH3 foci (Late G2) (Control: n=150; t1EK2/p53DN: n=150) or pan-nuclear staining (M-phase) (Control: n=150; t1EK2/p53DN: n=150) at 1.5 dpt (see Fig. S1a, b for protocol details). Anaphase onset was blocked with MG132. Control neurons were never positive for pH3. Three independent experiments were carried out. As control neurons never were never positive for pH3, statistical analysis was performed by comparing pH3-positive t1EK2 neurons with zero (***p<0.001, *p<0.05; one-tailed *t* test). **b** Percentage of LacZ/p53DN- (Control) or t1EK2/p53DN-expressing neurons with more than 5 γ-H2AX foci (Control: n=150; t1EK2/p53DN: n=150) at 1.5 dpt (see Fig. S1c for additional γ-H2AX results). Three independent experiments were carried out (***p<0.001; Fisher’s exact test). **c** Confocal image of pan-nuclear γ-H2AX staining of an interphase nucleus (left neuron) and γ-H2AX foci of a prophase-like nucleus (right neuron) (see also Video 1). **d** Percentage of LacZ/p53DN- (Control) or t1EK2/p53DN expressing neurons (Control: n=150; t1EK2/p53DN: n=150) at 1.5 dpt in late G2 or in M-phase after G2 checkpoint abrogation with MK1775 (see Fig. S1a, d for protocol details). Anaphase onset was blocked with MG132. Control neurons are never positive for pH3. Three independent experiments were carried out. As control neurons never were never positive for pH3, statistical analysis was performed by comparing pH3-positive t1EK2/p53DN neurons with zero (***p<0.001, *p<0.05; one-tailed *t* test). **e-g** Confocal images and 3D reconstructions of a pH3 foci-positive neuron consistent with late G2 (**e**) and neurons with pan-nuclear pH3 staining consistent with prometaphase/metaphase (**f**, see also Video 2, **g**). See also Fig. S1e-k. Blue, red and green arrows represent 3D orientation. **h** Confocal images and 3D reconstruction showing diffuse anillin staining in a t1EK2/p53DN-expressing neuron treated with MK1775 in prometaphase. Blue, red and green arrows represent 3D orientation. **i** Confocal images of a neuron with α-tubulin staining of putative mitotic spindle. Bottom panels: projection of slices 1 to 23 out of 50 to eliminate background cytoskeleton α-tubulin staining (see also Fig. S1l, m). Graphs represent mean ± s.e.m. Arrows identify RFP positive neurons. Scale bars: 25 μm (**c, e-i**), 10 μm (**i**, bottom panels).

In mitotic cells, G2/M transition can be blocked by G2 checkpoint activation^19^, which is triggered by DNA damage acquired during a genotoxic S-phase. Deregulated Cyclin E can induce DNA damage resulting from the aberrant activation of the DNA replication machinery^43–46^. Thus, we assessed whether DNA damage in t1EK2/p53DN neurons was present and could account for the overall block in G2/M transition. DNA damage was assessed by Phospho-Ser139 Histone H2AX (γ-H2AX) immunostaining^47^. We quantified the amount of neurons that were negative for γ-H2AX, had 1-5, 6-10, or more than 10 γ-H2AX foci, or showed γ-H2AX pan-nuclear staining. At 1.5 dpt, t1EK2/p53DN-expressing neurons generally presented more than 5 γ-H2AX foci and control neurons 5 or less foci (Fig. S1c). Thus, we analyzed whether the proportion of t1EK2/p53DN-expressing neurons with more than 5 γ-H2AX foci was significantly higher than that of control neurons. This analysis indicated that the proportion of neurons presenting more than 5 γ-H2AX foci was significantly higher in the t1EK2/p53DN-expressing group (78 %) than in the p53DN-expressing controls (24 %) (p<0.001, Fisher’s exact test) (Fig. 4b). Neurons with pan-nuclear γ-H2AX staining generally displayed interphase nuclei (Fig. 4c left neuron; for 360° view of DAPI-stained nuclei see Video 1). Neurons presenting reduced γ-H2AX foci formation could be occasionally found with prophasic-like mitotic chromatin condensation (Fig. 4c right; Video 1). These results support the possibility that DNA damage in neurons could be blocking M-phase entry by activating the canonical G2 checkpoint.

M-phase block exerted by G2 checkpoint signaling is reversible^19^. This checkpoint largely relies on Wee1 kinase inhibition of Cyclin B1/Cdk1 activity. In mitotic cells, G2/M transition can be induced by inhibiting Wee1 kinase with MK1775^48^. To test whether Wee1 kinase was preventing neurons from entering M-phase, we abrogated G2 checkpoint signaling by inhibiting Wee1 and assessed pH3 immunostaining. Again, we blocked anaphase onset with MG132^42^ to capture all neurons that entered M-phase (see Fig. S1a, d for protocol details). Control MAP2-positive neurons were not found in late G2 or M-phase (n=150) (Fig. 4d). In contrast, Wee1 inhibition with MK1775 resulted in a substantial increase of t1EK2/p53DN-expressing neurons in M-phase (31.3 %) (*t*_149_=8.246, p<0.001) (Fig. 4d, f, g, Fig. S1f-k; for 360° view of RFP/pH3 labeling illustrated in Fig. 4f, see Video 2). In addition, 37.3 % (*t*_149_=9.422, p<0.001) of t1EK2/p53DN-expressing neurons were still found in late G2 as reflected by pH3 foci (Fig. 4d, e, Fig. S1e). The total percentage of neurons in late G2 and M-phase after Wee1 inhibition (68.6 %) was consistent with the percent of BrdU positive neurons at 2 (61.2 %) or 3 dpt (61.8 %) (Fig. 3a).

Neuronal M-phase was further confirmed by anillin immunolabeling, which shuttles from the nucleus to the cell cortex during the prophase to prometaphase transition^49^. In neurons with nuclei displaying characteristics of prometaphase/metaphase, anillin immunostaining was released from the cell nucleus (Fig. 4h). We also performed α-tubulin immunostaining of the mitotic spindle of neurons in M-phase. Although α-tubulin staining resulted in extensive background signal due to its presence in the cytoskeleton, putative mitotic spindles with intense α-tubulin labeling could be identified in neurons with chromatin condensation, consistent with prometaphase/metaphase (Fig. 4i, Fig. S1l, m). Finally, we performed time-lapse experiments to follow live neurons into M-phase. We expressed Histone H2B tagged with EGFP (H2B-GFP) to follow DNA dynamics and RFP reporter protein to study morphology. H2B-EGFP signal was consistent with mitotic chromatin condensation (Video 3, right). Chromatin condensation was usually accompanied by a loss of RFP signal from dendrites. This may reflect a thinning and or retraction of the dendrites that is coordinated with M-phase entry. In contrast, t1EK2/p53DN expressing neurons in interphase remained with unaltered gross morphology (Video 3, left). In summary, primary neurons can progress through S and G2-phases and enter M-phase. M-phase entry in neurons is gated by G2 cell cycle checkpoint signaling as in mitotic cells.

### Differentiated neurons can undergo cytokinesis

We again performed time-lapse experiments to see whether neurons can undergo cytokinesis. To improve the chances of cytokinesis, we used caffeine to inhibit G2 checkpoint signaling^50^ upstream of Wee1^19^ and co-expressed Topoisomerase-2α (TOP2α) to aid in the decatenation of intertwined sister chromatids^51^ (see Fig. S2a for protocol details). Although sparse and highly asynchronous, attempts at anaphase/cytokinesis could be detected in 12.9 % (n=93) of recorded neurons. Neurons largely underwent anaphase/cytokinesis failure (n=8) (Video 4). Despite this, some neurons were able to complete cell division (n=4). One of the daughter neurons could die whilst the other remained viable (Fig. 5a, b; Video 5; see Fig. S2a, b for protocol details), evidencing that neuronal cell division was complete. Alternatively, both neurons could remain viable for hours after abscission (Fig. 5c, d; Video 6; see Fig. S2a, c for protocol details).

**Fig. 5.**
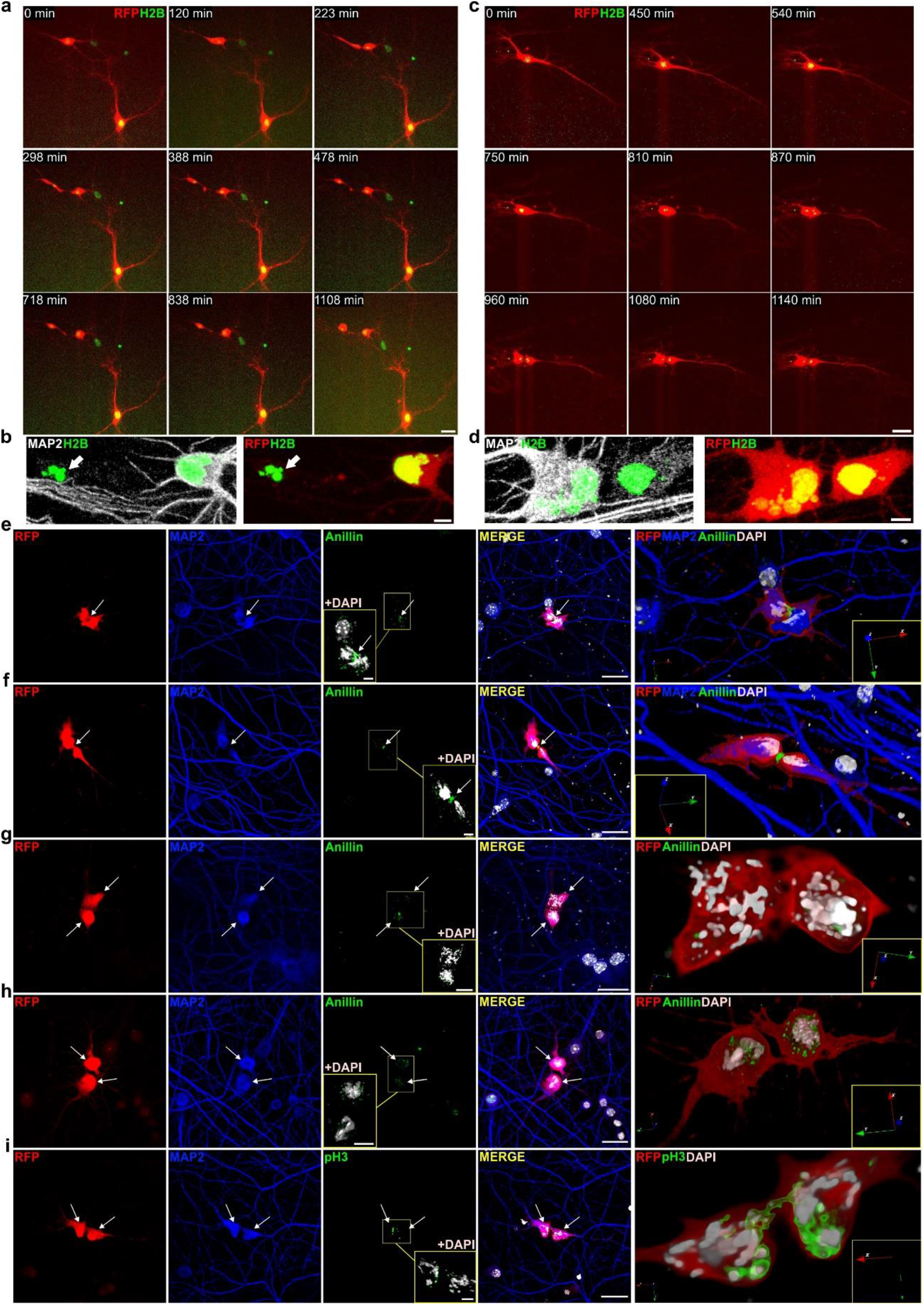
Differentiated neurons can undergo cytokinesis. **a** Time-lapse video frames of a neuron undergoing cytokinesis (Video 6). There is a delay between the formation of the intercellular bridge and final abscission, suggestive of the presence of a chromatin bridge. **b** Confocal images of daughter neurons (evidenced by MAP2 immunostaining) in **a**, confirming that abscission was complete. One daughter neuron survived (right) while the other (left) displayed pyknotic nucleus (white arrow). **c** Time-lapse video frames of a neuron undergoing cytokinesis (Video 7). **d** Confocal images of daughter neurons (evidenced by MAP2 immunostaining) in **c**, confirming that abscission was complete. The daughter neuron on the left is multinucleated, indicative of aneuploidy. **e-h** Confocal images of anillin immunolabelling. Anillin stains the cleavage furrow/midbody during neuronal cytokinesis (**e**, **f;** for 360° view of **f** see Video 7). White arrows indicate intercellular bridge/midbody. See also Fig. S2f-h. Anillin is recycled from the midbody in putative daughter neurons after completing abscission (**g**, **h**). **i** Confocal images of a pH3 positive neuron undergoing cytokinesis (for 360° view see Video 8). The intercellular bridge is traversed by pH3 positive chromatin, evidencing the presence of a chromatin bridge. White arrows identify daughter neurons (**g-i**). Blue, red and green arrows represent 3D orientation (**e-i**). H2B: Histone H2B tagged with EGFP. Scale bars: 25 μm (**a**, **c**, RFP and H2B-EGFP stills), 5 μm (**a**, **b**, bottom panels), 25 μm (**e-i**).

We also assessed the assembly of the cytokinesis machinery by anillin immunocytochemistry. Anillin localizes to the cleavage furrow at cytokinesis onset and remains in the intercellular bridge after cleavage furrow ingression^49^. The detection of cytokinesis by immunocytochemistry only captures neurons undergoing cytokinesis at a single time-point. However, cytokinesis onset in different neurons could be hours apart (Fig. 5a, c; Videos 5, 6). To achieve some degree of synchronization, we forced anaphase/cytokinesis onset by abrogating the Spindle Assembly Checkpoint (SAC) with the Mps1 inhibitor AZ3146^52^ (see Fig. S2d, e for protocol details). Confirming our time-lapse experiments, neurons assembled the cytokinesis machinery (Fig. 5e, f; Fig. S2f-h). We found that, in most cases, cleavage furrow ingression began (Fig. S2f) and proceeded asymmetrically (Fig. S2g, h) up to the formation of the intercellular bridge/midbody (Fig. 5e, f; for 360° view of Fig. 5f see Video 7). We were also able to detect putative daughter neurons after abscission completion (Fig. 5g, h). We also performed pH3 immunolabelling to characterize chromatin decondensation during anaphase/telophase. The loss of pH3 staining begins at anaphase and is completed prior to detectable chromosome decondensation in telophase^41^. In agreement, neurons locked in cytokinesis by chromatin bridges displayed both patterns of staining (Fig. 5i; for 360° view of Fig. 5i, see Video 8). This suggests that the pH3-positive chromatin trapped in the intercellular bridge was still condensed while the portion of the chromatin outside the intercellular bridge lacking pH3 staining was already decondensed in a telophase-like state.

In sum, we conclude that cell cycle regulation in S, G2, and M phases in neurons is shared with mitotic cells.

### Neurons recover AIS integrity after M-phase exit without cell division

Neuronal cell cycle re-entry in neuron disease is associated to cell death instead of cell division^15,16^. To determine whether cell cycle re-entry without cell division is necessarily fatal, we assessed the viability of t1EK2/p53DN/TOP2α-expressing multinucleated neurons, which were frequently detected in our cultures. Multinucleation requires M-phase entry and, therefore, enabled the identification of neurons that had entered and exited the cell cycle without cell division. As a control, we did not detect any evidence of multinucleation in controls transfected neurons in BrdU (n=491, Fig. 1a; n=1,043, Fig. 3a) or pH3 (n=150, Fig. 4a; n=150, Fig. 4d) experiments. As neurons expressed the p53DN mutant, we assessed AIS integrity rather than active caspase-3 to study viability. The AIS sustains neuronal polarity and integrates synaptic input to generate action potentials^53,54^, and previous studies demonstrate that AIS integrity is required to maintain neuronal viability^54,55^. Within the AIS, AnkyrinG (AnkG) is the main structural constituent, necessary to maintain AIS functions and neuronal integrity^53^. We analyzed AnkG dynamics in t1EK2/p53DN/TOP2α-expressing neurons in interphase, prophase, and prometaphase/telophase 3 h after G2 checkpoint abrogation. In multinucleated neurons we assessed AnkG 6 and 9 h after abrogating the G2 checkpoint. In control LacZ/p53DN/TOP2α-expressing neurons AnkG was assessed 9 h after G2 checkpoint abrogation.

AnkG signal varied significantly across cell cycle stages [Welch’s F test (5; 20,479) = 11.617, p<0.001] (Fig. 6a, b), evidencing that AIS integrity fluctuated during the cell cycle. Multiple comparisons using the Games-Howell post-hoc test indicated that the AnkG signal of neurons in prophase was not significantly different from that of neurons in interphase (p=0.448) or control transfected neurons (p=0.703) (Fig. 6a, b). However, the AnkG signal did decrease significantly in prometaphase to telophase when compared to interphase (p=0.003), prophase (p=0.023) or control neurons (p=0.009). These results evidence a loss of AIS integrity in M-phase, in between prometaphase and telophase. 6 h after inducing G2/M transition, multinucleated neurons had still not recovered the AnkG signal when compared to interphase (p=0.033), prophase (p=0.033), or control neurons (p=0.041). However, 9 h after inducing G2/M transition, multinucleated neurons significantly increased their AnkG signal with respect to neurons in between prometaphase and telophase (p=0.005), achieving an AnkG signal that was non-significantly different from control (p=0.957), interphase (p=0.999), or prophase (p=0.349) neurons. The recovery of AnkG signal in multinucleated neurons was gradual, as multinucleated neurons 9 h after inducing G2/M transition displayed a non-significant trend to increase the AnkG signal when compared to 6 h (p=0.065). Therefore, the AIS is coordinated with the neuronal cell cycle. After cell cycle exit without cell division, neurons can progressively yet fully recover the AIS.

**Fig. 6.**
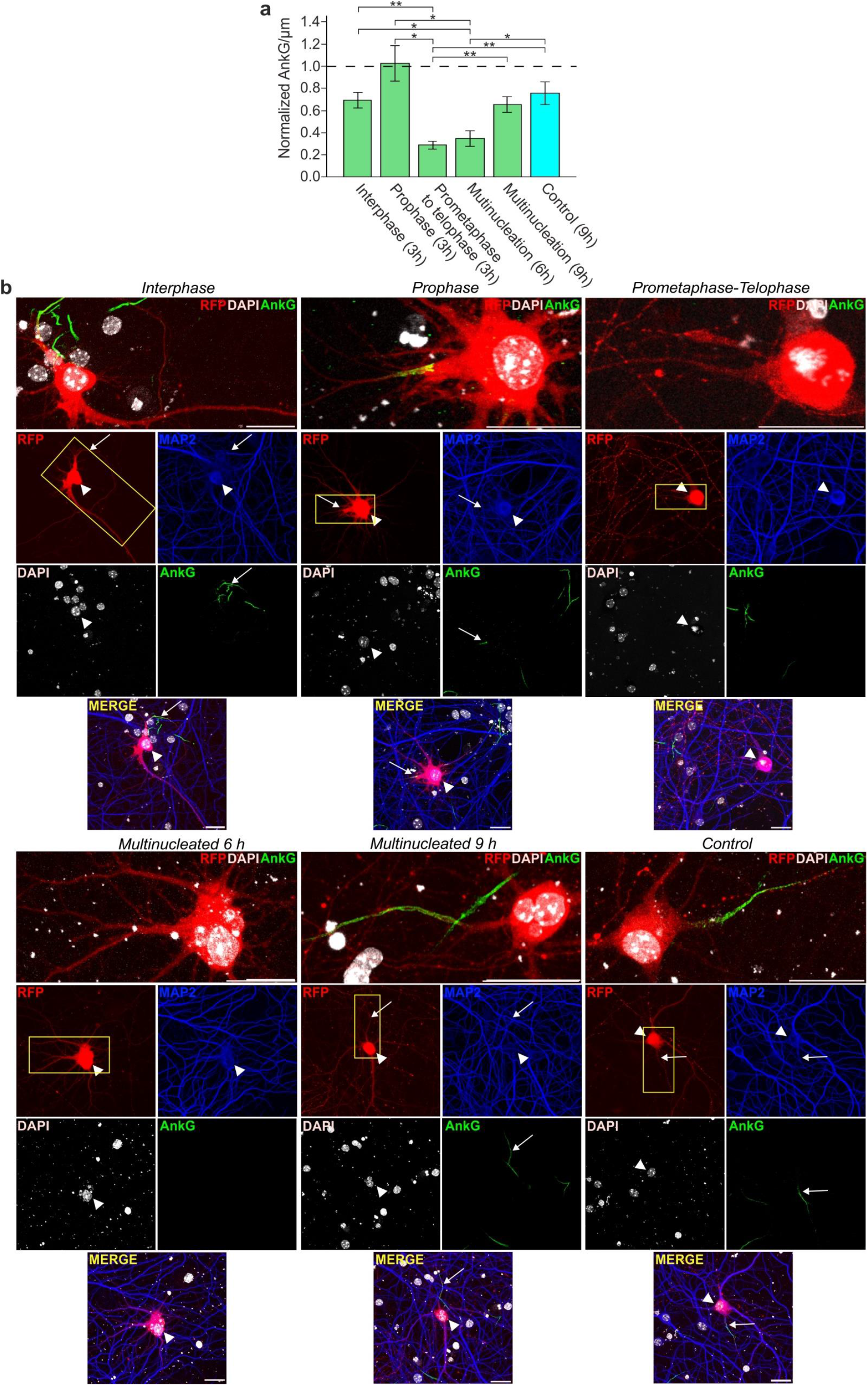
Neurons recover AIS integrity after M-phase exit without cell division. **a** AnkG signal per μm in t1EK2/p53DN/TOP2α-expressing neurons in different stages of the cell cycle (green) and in control neurons expressing LacZ/p53DN/TOP2α (blue) at the indicated time points after suppressing the G2 checkpoint with MK1775, normalized to neighboring RFP-negative/MAP2-positive neurons. All groups were treated with AZ3146 to abrogate the SAC 2.5 h after MK1775 treatment. A single experiment was carried out (**p<0.01, *p<0.05; Welch’s F test and two-sided Games-Howell post-hoc tests; Interphase: n=9, Prophase: n=7, Prometaphase to telophase: n=9, Multinucleation 6 h: n=8, Multinucleation 9 h: n=10, Control: n=10). **b** Confocal images of AnkG-immunolabelled neurons in interphase, prophase, and prometaphase to telophase, multinucleated neurons 6 and 9 h post transfection, and control neurons. High magnification of AIS (boxes) is shown for each case. Arrowheads identify RFP positive neurons and arrows their respective AIS when present. Graphs represent mean ± s.e.m. Scale bar: 25 μm.

## Discussion

It was originally believed that the postmitotic status of neurons prohibited any form of cell cycle activity. This view is now outdated. There is ample evidence of neuronal cell cycle re-entry in human pathologies such as AD, PD, AML or brain injury, as well as in animal models and in primary neuronal cultures^15,16^. Notwithstanding, neuronal cell cycle re-entry in differentiated mature neurons does not result in cell division but it is instead associated to increased susceptibility to cell death^16^. Congruently, the activation of the cell cycle machinery itself has been deemed pathological^17^. Therefore, it is generally thought that the regulation and deregulation of the cell cycle machinery in neurons differs from that of mitotic cells. We provide evidence to the contrary. Primary neurons undergo apoptosis upon cell cycle re-entry. As with mitotic cells, apoptosis in S-phase can be prevented by suppressing p53 signaling. Rescued neurons progress through S-phase into G2, where they are arrested at the G2/M transition. Inhibition of G2 checkpoint kinase Wee1 reveals neurons readily enter M-phase and a subset of neurons can undergo cytokinesis. Therefore, although our results have to be confirmed in vivo, we have shown that a large part of the regulation of the cell cycle in S, G2 and M-phase in primary neurons at post-synaptogenesis stages of development is shared with mitotic cells. Hence, cell cycle-induced cell death, lack of M-phase entry, or lack of cell division in neurons can be accounted for by standard cell cycle regulation present in mitotic cells. We have also shown that neurons that exit the cell cycle without undergoing cell division, thus becoming multinucleated, can recover the AIS, which is necessary to sustain neuronal viability^54,55^.

Viable cell cycle exit is consistent with reports showing that neurons that re-enter the cell cycle in AD can remain viable for months to years^16,56^. Furthermore, the existence of multinucleated neurons in AD suggest that at least some neurons that re-enter the cell cycle can reach M-phase^57^. However, in current models, cell cycle re-entry invariantly results in cell death. This prevents understanding how neurons can survive cell cycle re-entry in neurodegenerative diseases and underscores the need for models of viable neuronal cell cycle entry^16^. In this study, we provide a model in which cell cycle-related cell death is prevented, and that allows the study of dividing or multinucleated neurons. We propose that canonical cell cycle regulation in neurons will afford a new framework to study cell cycle deregulation in neurodegeneration and basic research.

## Methods

### Plasmids

Construction of a vector encoding a truncated form of Cyclin E (Trunk 1), which lacks the first 39 amino acids of Cyclin E^23^, fused to Cdk2 isoform 1 (NCBI accession number NP_001789) (t1EK2) was based on the EK2 fusion protein described previously^58^. Briefly, a synthetic DNA construct encoding for t1EK2 was firstly cloned into the pUC vector, then subcloned into the HindIII and XbaI sites of pcDNA3 vector (ThermoFisher Scientific) and finally sequence verified. The pcDNA6/V5-His/lacZ vector expressing LacZ was purchased from Invitrogen. The pRFPRNAiC vector expressing red fluorescent protein (RFP), provided by Stuart Wilson (University of Sheffield, UK), has been previously described^59^. The T7-p53DD-pcDNA3 vector expressing a dominant negative form of p53^40^ was a gift from William Kaelin (Addgene plasmid # 25989). The H2B-GFP vector expressing a Histone-GFP fusion protein^60^ was a gift from Geoff Wahl (Addgene plasmid #11680). The TOP2α-WT pcDNA3.1(+) vector expressing topoisomerase IIα^61^ was a kind gift from Corrado Santocanale (National University of Ireland).

### Antibodies and inhibitory compounds

Primary antibodies: The anti-BrdU rat monoclonal antibody (mAb) [BU1/75 (ICR1)] (AbDSerotec) was used at 1:200 dilution. The anti-RFP rabbit polyclonal antibody ab62341 (Abcam) was used at 1:100 dilution. The anti-MAP2 chicken antibody ab5392 (Abcam) was diluted 1:5000-1:12000. The cleaved Caspase-3 rabbit antibody (Cell Signaling Technology) was used at 1:400 dilution. The anti-phospho Histone H3 (ser10) rabbit polyclonal antibody 06-570 (Millipore) was used at 1:500 dilution. The anti-anillin rabbit polyclonal antibody ab154337 (Abcam) was used at 1:100 dilution. The anti-α-tubulin mouse monoclonal antibody [DM1A] ab2791 (Abcam) was used at 1:10000 dilution. The anti-phosphoH2AX (ser139) mouse monoclonal antibody [9F3] ab26350 (Abcam) was used at 1:500 dilution. The anti-Ankyrin-G mouse antibody [N106/36] (NeuroMab) was diluted at 1:150. Secondary antibodies: Alexa Fluor 488 Goat Anti-Rat IgG (H+L) (Life Technologies), Alexa Fluor 488 Goat Anti-Mouse IgG (H+L) (Life Technologies), Alexa Fluor 488 Goat anti-Mouse IgG2a A-21131 (Invitrogen), Alexa Fluor 488 Affinipure Goat Anti-Rabbit IgG (H+L) (Jackson Immunoresearch), Alexa Fluor 594 Goat Anti-Rabbit IgG (H+L) (Life Technologies), and Alexa Flour 647 Goat Anti-Chicken IgY (H+L) A-21449 (Invitrogen) were used at 1:1000 dilution. Inhibitors: Cdk1 inhibitor RO3306 (ref. 1530) was used at 9 μM, Wee1 inhibitor MK1775 (ref. 1494), used at 900 nM, and Mps1 inhibitor AZ3146 (ref. 1642), used at 10 μM, were purchased from Axon Medchem. The Eg5 inhibitor monastrol (ab14087), used at 100 μM, and the proteasome inhibitor MG132 (ab141003), used at 10μM, were purchased from Abcam. Caffeine (Sigma), used at 3 mM, was diluted in Neurobasal without L-Glutamine (ThermoFisher Scientific) and filter sterilized with 0.2 μm syringe filters (Acrodisc).

### Hippocampal Cultures

10-mm diameter coverslips (Menzel Glässer) were placed in 65% nitric acid (Merck) overnight. Coverslips were then washed 4 times in Milli-Q water (Millipore), once in pure ethanol (Merck) and air-dried. Coverslips were sterilized overnight in oven at 185°C, placed in CELLSTAR cell culture dishes with 4 inner rings (Greiner bio-one) and UV sterilized for 1 h. Coverslips were coated with 0.5 mg/ml polyornithine (PLO) (Sigma-Aldrich) (prepared in 150 mM borate buffer, pH8.4) for at least 24 h. PLO was washed twice in sterilized Milli-Q water. CELLSTAR culture dishes were left in a humidified atmosphere containing 5 % CO2 at 37°C with 2ml of neuronal plating medium consistent of Dulbecco’s modified Eagle medium (DMEM) with D-Glucose, L-Glutamine and Pyruvate (ThermoFisher Scientific) containing 10 % fetal bovine serum (FBS) (Life technologies) and penicillin-streptomycin (25 U/ml) (Gibco). Primary hippocampal neurons derived from CD-1 mice (Charles River) were harvested from embryonic day 17, staged as previously described^62^. Pups were decapitated and brains were placed in cold Hanks’ balanced salt solution (HBSS) without calcium chloride, magnesium chloride, nor magnesium sulphate (Gibco). Hippocampi were then dissected and incubated for 15 min at 37°C in 5 ml HBSS containing 2.5 mg/ml trypsin (Gibco) in a 15 ml conical centrifuge tube. 1 mg/ml DNAse (Roche) was added for the last 30 s. Hippocampi were then washed 5 times in 5 ml of HBSS at 37°C each time. Mechanical dissociation followed in 1ml of HBSS at 37°C by passing hippocampi through a Pasteur pipette 10-15 times. Non-dissociated tissue was allowed to collect at the bottom by gravity and clean supernatant with dissociated cells transferred to a new 15 ml centrifuge tube. 1 ml of HBSS at 37°C was added to the remaining tissue and dissociated again with a flame polished Pasteur pipette to reduce the diameter of the tip 10-15 times. Non-dissociated tissue was allowed to collect at the bottom by gravity. Clean supernatant with dissociated cells was transferred to the same centrifuge tube in which cells from the first dissociation were collected. Neurons were transferred at a density of 24.000 cells/cm^2^ to CELLSTAR cell culture dishes (Greiner bio-one) with neuronal plating medium. Neurons were allowed to attach to coverslips for 3 to 4 h in a humidified atmosphere containing 5 % CO2 at 37°C. Once neurons were settled onto the coverslips, neuronal plating medium was washed with maintenance medium consistent of Neurobasal without L-Glutamine (ThermoFisher Scientific) medium supplemented with B27 (ThermoFisher Scientific), penicillin-streptomycin (25 U/ml) (Gibco) and GlutaMAX (Gibco) and differentiated with the same medium up to 3-4 DIV. Half of the culture medium was exchanged by fresh maintenance medium without penicillin-streptomycin every 3-4 days.

### Lipofection

All lipofections were carried out in hippocampal neurons differentiated for 15 DIV. Two or three days prior to lipofection, half of the culture medium was changed for fresh growth medium without penicillin-streptomycin. In order to increase viability, recovery medium was prepared from conditioned medium of neuronal cultures and used after lipofection. Thus, 30-60 min prior to lipofection, half of the conditioned medium of each neuronal culture (1 ml) was removed and collected into a clean culture dish containing 1 ml of fresh growth medium. BrdU (5 μg/ml) and or inhibitors were added to the recovery medium as needed. The recovery medium was then incubated in a humidified atmosphere containing 5 % CO_2_ at 37°C for the duration of the lipofection. Neurobasal medium without supplements pre-warmed to 37°C was added to each of the neuronal cultures. Cultures were returned to the incubator for a minimum of 30 min to stabilize CO_2_ levels. To maximize viability, neurons were lipofected in their own culture dish and remaining conditioned medium, which was free of penicillin-streptomycin by 15 DIV. Neurons were transfected with Lipofectamine 2000 (Invitrogen). Lipoplexes were prepared at a ratio of 1 μl of Lipofectamine per 1 μg of DNA. For immunocytochemistry experiments, RFP DNA was always transfected at a 1:20 ratio. Total transfected DNA per culture dish for t1EK2/RFP and LacZ/RFP experiments was 6.4 μg and total transfected DNA per culture dish for t1EK2/p53DN/RFP, LacZ/p53DN/RFP, t1EK2/p53DN/TOP2α/RFP, LacZ/p53DN/TOP2α-RFP experiments was 12.85 μg. For time-lapse experiments, RFP DNA was also transfected at 1:20. Total transfected DNA for t1EK2/p53DN/H2B-EGFP was 10.27 μg and for t1EK2/p53DN/TOP2α/H2B-EGFP it was 12.44 μg. Lipofectamine was mixed with Neurobasal to a final volume of 250 μl per culture dishand left for 3-5 min. Next, plasmid DNA diluted in 250μl of Neurobasal per culture dish was added and left for 20 min at room temperature. Thereafter, 0.5 ml of the mix was added to each culture dish and left for 90 min. Lipofectamine was then washed out 3 times with 2 ml of growth medium medium pre-warmed to 37°C. After washout, the recovery medium was added to its corresponding neuronal culture. For cytokinesis experiments, cultures were supplemented with 1xEmbryoMax Nucleosides (Merck)^45^.

### Immunocytochemistry

Hippocampal cultures were fixed for 15 min with 4 % paraformaldehyde (PFA) at RT, and permeabilized for 30 min with phosphate buffered saline (PBS) containing 0.05 % Triton X-100 (Sigma-Aldrich) (0.1% for AnkG staining) (PBTx). For BrdU inmunolabeling, DNA was denatured for 30 min at RT with 2N HCl/0.33× PBS, and then neutralized with three 15-min washes with 0.1 M sodium borate, pH 8.9, and a wash of 5 min with PBTx. Cultures were then incubated for 30 min at RT with PBTx containing 10 % FBS to block antibody unspecific binding, followed by a 1-h incubation at RT with PBTx/1 % FBS and the appropriate primary antibodies. After 4 washes in PBTx, cultures were incubated for 1 h at RT in PBTx containing 1 % FBS and the appropriate secondary antibodies. After 4 additional washes as above, DNA labelling was performed using PBS containg 100 ng/ml 4′,6-diamidino-2-phenylindole (DAPI) (Sigma-Aldrich) and the preparations were mounted in glycerol (Panreac)/PBS (1:1).

### Image analysis and cell counting studies

For BrdU incorporation time-course experiments, the proportion of BrdU positive neurons out of all MAP2 positive transfected neurons of 10-mm diameter coverslips (MenzelGlässer) was calculated in 3 independent experiments. To assess apoptosis, the proportion of active Caspase-3^38^ positive neurons of all MAP2 positive transfected neurons was calculated in each of 3 independent experiments. DNA damage was assessed with Phospho-Ser139 Histone H2AX (γ-H2AX)^47^. The proportion of neurons with more than 5 γ-H2AX foci or γ-H2AX pan-nuclear staining out of all MAP2 positive transfected neurons was calculated in each of 3 independent experiments. Phospho-Ser10 Histone H3 (pH3) immunostaining was used to assess late G2 and M-phase entry^42^, wherein the proportion of neurons positive for foci (late G2) or pan-nuclear (M-phase) staining out of all MAP2 positive transfected neurons was calculated in each of 3 independent experiments in each treatment condition. Cell counting in BrdU, active Caspase-3, pH3 and γ-H2AX experiments was done with 20x (Zoom, 1.5) or 40x objectives with AF 6500-7000 fluorescence microscope (Leica) and a DU8285 VP-4324 camera (Andor). Confocal microscope images were acquired with 40x or 63x objectives using upright or inverted TCS SP5 confocal microscopes (Leica). For three-dimensional (3D) image reconstructions, images were taken with a 63x objective with 1.7- to 3.6-times zoom, in 50 to 69 z-stacks of 0.13 to 0.25 μm step-sizes. 3D reconstructions and rotating 3D videos were generated using Icy bioimage informatics platform^63^.

### Live imaging of neuronal cell cycle

Neurons were cultured as described above in μ-Dish 35mm, high Grid-500 cell culture dishes with ibidi polymer coverslip bottom (Ibidi) coated with 0.5 mg/ml polyornithine (Sigma-Aldrich). Live neuronal imaging was performed in a sealed chamber at 37°C and 5 % CO2 with 20x objective of a AF 6500-7000 fluorescence microscope and recorded with DU8285 VP-4324 camera. Pictures and videos were generated using Leica Application Suite X (Leica). Time intervals between pictures and experiments duration are described in each video. For cell counting studies of anaphase and cytokinesis, neurons were identified by morphological criteria based on RFP signal. When in doubt, cells were either not included in the quantification or neuronal identity confirmed by MAP2 immunostaining and included in the quantification. Binucleated neurons were not quantified. Neurons that presented obvious signs of cell death during the first hour of recording were not included in the quantification. Chromosome segregation during anaphase was identified by H2B-GFP^60^ and cleavage furrow ingression by RFP.

### Measurement of the AIS

AnkyrinG (AnkG) was used to evaluate changes in the AIS of neurons at different stages of the cell cycle. All groups were treated with Wee1 inhibitor MK1775^49^ (900 nM) to abrogate the G2 checkpoint and with the Mps1 inhibitor AZ3146^52^ (10 μM) 2.5 h later to abrogate the SAC. Neurons in the interphase, prophase and prometaphase to telophase groups were fixed 3 h after G2 checkpoint abrogation. Stages in between prometaphase and telophase could not be reliably distinguished by nuclear morphology and were thus grouped together. Multinucleated neurons were used to assess premature M-phase exit and were fixed 6 and 9 h after G2 checkpoint abrogation. Multinucleated neurons still presenting mitotic chromatin condensation or evidence of pyknosis were not included in the quantification. Neurons in all of the aforementioned groups co-expressed t1EK2/p53DN/TOP2α/RFP. Control neurons expressing LacZ/p53DN/TOP2α/RFP were fixed 9 h after G2 checkpoint abrogation. Images were acquired on an upright Leica SP5 confocal microscope with a 63× objective, 1024 × 1024 pixels in z stacks with 0.5-μm steps and a Z-projection was obtained. Measurements of AnkG fluorescence intensity were performed by Fiji-ImageJ software. We drew a line starting at the limit of neuronal soma identified by MAP2 staining, and extended it along the AnkG staining or the RFP signal of the axon. Data were obtained after background subtraction. Then, data were smoothed every 1 μm using Sigma Plot 12.5 software. AIS start and end positions were obtained following the criteria described previously^64^. Then total fluorescence intensity for each AIS was obtained. Total fluorescent intensity was then divided by the length of the AIS to obtain the mean AnkG fluorescence per 1 μm (AnkG/μm). To avoid variability between cell cultures and treatment exposure times, we normalized the mean AnkG/μm of transfected neurons to the nearest neighboring non-transfected neurons in the same image.

### Statistical Analysis

Statistical analysis was performed using SPSS (version 24.0). Statistical analysis of BrdU time-course experiments of pH3 experiments were done with one-sample *t* test against 0 because control neurons never incorporated BrdU or were positive for pH3 (α=0.05, one-tailed). For BrdU time-course experiments, the mean percent BrdU incorporation of the treatment group was compared to 0 in each time-point. Analysis of active-caspase-3 and γ-H2AX experiments were done with Fisher’s exact test (α=0.05, two-tailed). For AnkG/μm in AIS experiments, outliers were identified by boxplots. Normality was assessed with Shapiro-Wilk test of normality and Normal Q-Q Plots. Homogeneity of variances with Levene’s test for equality of variances. Omnibus testing was performed with Welch’s F test and post-hoc multiple comparisons with the Games-Howell method (α=0.05, two-tailed). Quantitative data are expressed as mean ± standard error of the mean (s.e.m.). Significance was p<0.05 (*), p<0.01 (**), and p <0.001 (***).

### Data availability

The datasets generated during and/or analyzed during the current study are available from the corresponding author on reasonable request.

## Acknowledgements

We thank Stuart Wilson, William Kaelin, Geoff Wahl, and Corrado Santocanale for the plasmids used in this study, and S. Casas-Tintó, and F. Wandosell for the critical reading of the manuscript. We also thank the Department of Statistics of SGAI-CSIC. Funding: The study was supported by Ministerio de Economía y Competitividad grant SAF2015-68488-R (J.M.F.), SAF2015-65315-R (J.J.G.), and Ministerio de Educación, Cultura y Deporte grant FPU1305084 (C.C.W.).

## Author contributions

C.C.W., J.M.F., and J.J.G. conceived the study, C.C.W. performed the experiments and analyzed data, W.Z. analyzed data, I.P.-P. analyzed data, E.B.-A. provided expertise and feedback, C.C.W. and J.M.F. wrote the manuscript.

## Additional information

**Supplementary Information** accompanies this paper.

## Competing interests

The authors declare no competing interests.

---

**Fig. S1.**
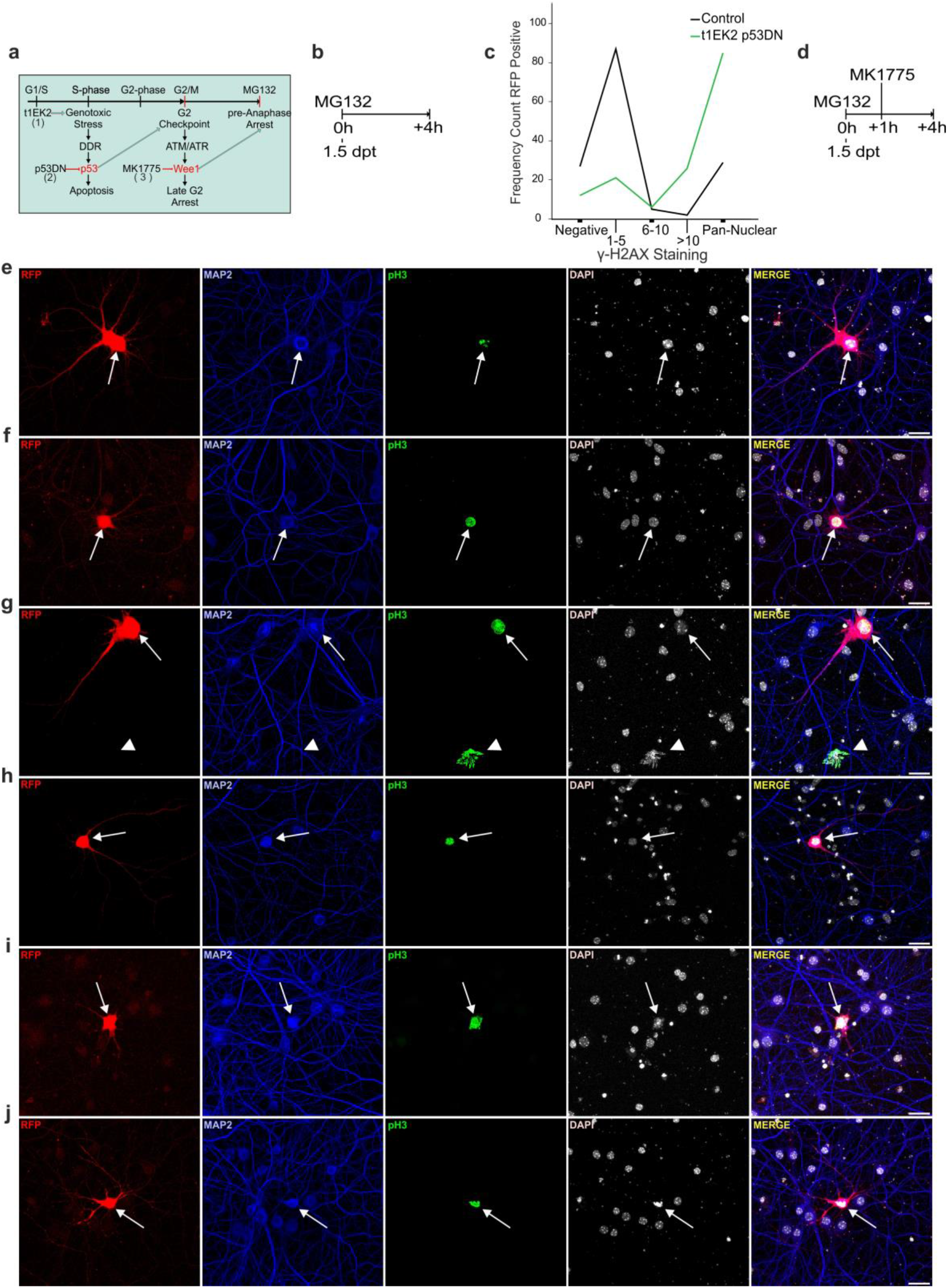
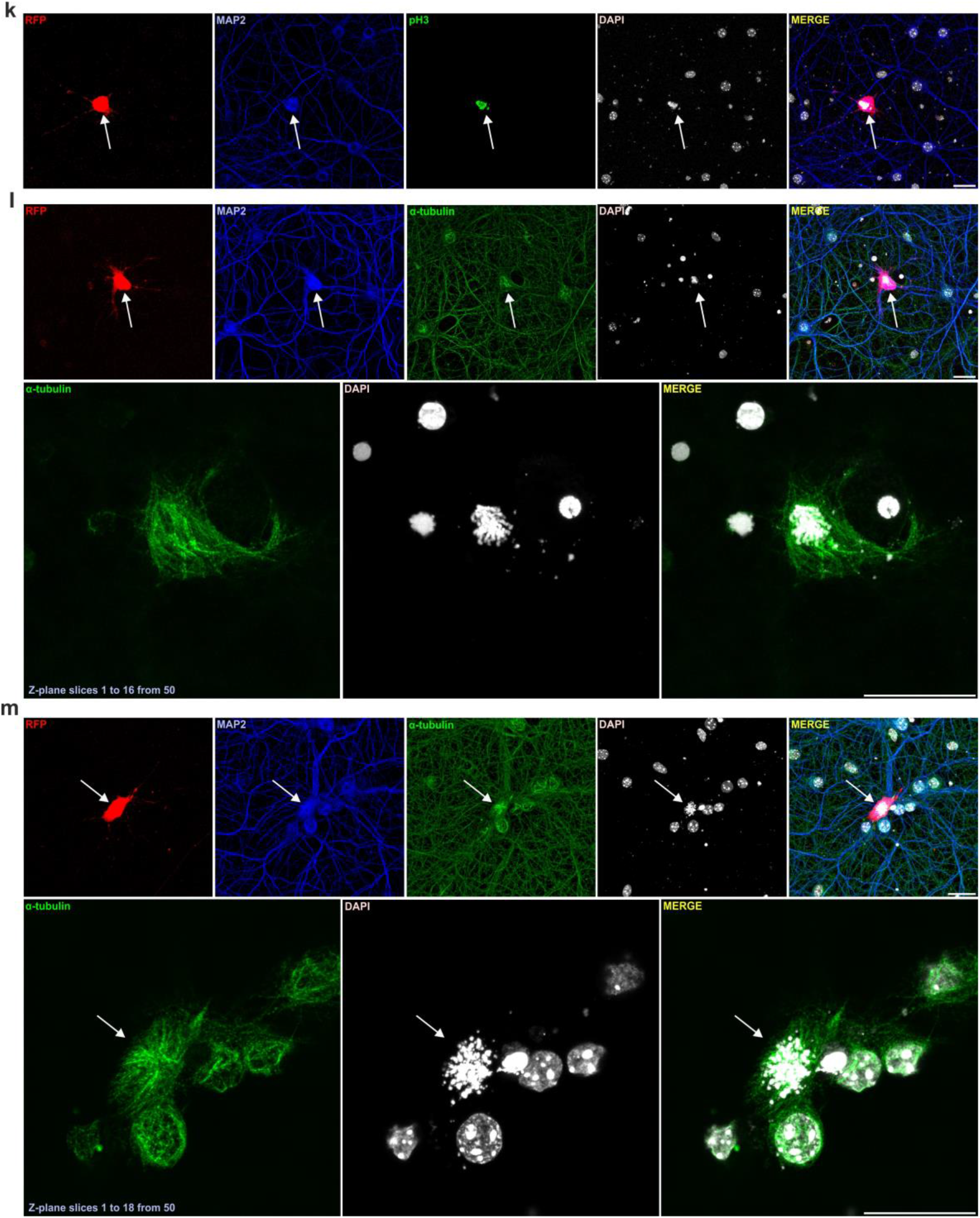
Wee1 inhibition enables G2/M transition in differentiated neurons. **a** Diagram depicting cell cycle regulation and manipulations performed in this study. t1EK2 induces G1/S transition (1) and DNA damage that can lead to p53-dependent apoptosis^18^. In turn, the loss of function of p53 affords viability and cell cycle progression (2). Due to dysfunctional G1/S and S checkpoint regulation, p53-deficient cells rely heavily on G2 checkpoint arrest to amend DNA damage prior to M-phase entry^19^. The G2 checkpoint relies on Wee1 kinase. Wee 1 inhibition with MK1775 (3) can abrogate the G2 checkpoint and afford M-phase entry^48^. **b** Experimental protocol to assess neurons in late G2 and M-phases without suppressing G2 checkpoint signaling. At 1.5 dtp, the proteasome was inhibited with MG132 (10μM) to prevent mitotic exit^42^ and the accompanying loss of pH3 staining^41^. Neurons were fixed for immunocytochemistry 4 h later. **c** DNA damage assessed by γ-H2AX. Frequency of LacZ/p53DN (Control, n=150) and t1EK2/p53DN (n=150) expressing neurons negative for γ-H2AX, with 1 to 5, 6 to 10, or above 10 γ-H2AX foci, or pan-nuclearγ-H2AX staining. Neurons distributed to the negative to 5 γ-H2AX foci groups or to the more than 5 γ-H2AX groups. Therefore, analysis of DNA damage was assessed by comparing the percentage of neurons bearing more than 5 γ-H2AX foci between t1EK2/p53DN and LacZ/p53DN groups (Fig. 4b). **d** G2 checkpoint abrogation protocol to assess G2/M transition. At 1.5 dtp, the proteasome was inhibited with MG132 (10μM) to prevent mitotic exit^42^ and the accompanying loss of pH3 staining^41^. One hour later, MK1775 (900 nM) was added to inhibit Wee1 and abrogate G2 checkpoint signaling. Neurons were fixed for immunocytochemistry 4 h after addition of MG132. **e-k** Confocal images of a neuron in late G2 positive for pH3 foci (**e**), pan-nuclear pH3 staining with chromatin condensation consistent with prophase (**f, g, h**), prometaphase (**i**) and prometaphase/metaphase (**j, k**). **l, m** Confocal images of neuron with α-tubulin immunostaining of putative mitotic spindles. Bottom panels: projection of slices 1 to 16 (**l**) and 1 to 18 (**m**) out of 50 to eliminate background, cytoskeletal α-tubulin staining. Arrows identify RFP positive neurons and arrowheads MAP2-negative glial cells. Scale bars: 25 μm (**e-m**), 10 μm (**l, m** bottom panels).

**Fig. S2.**
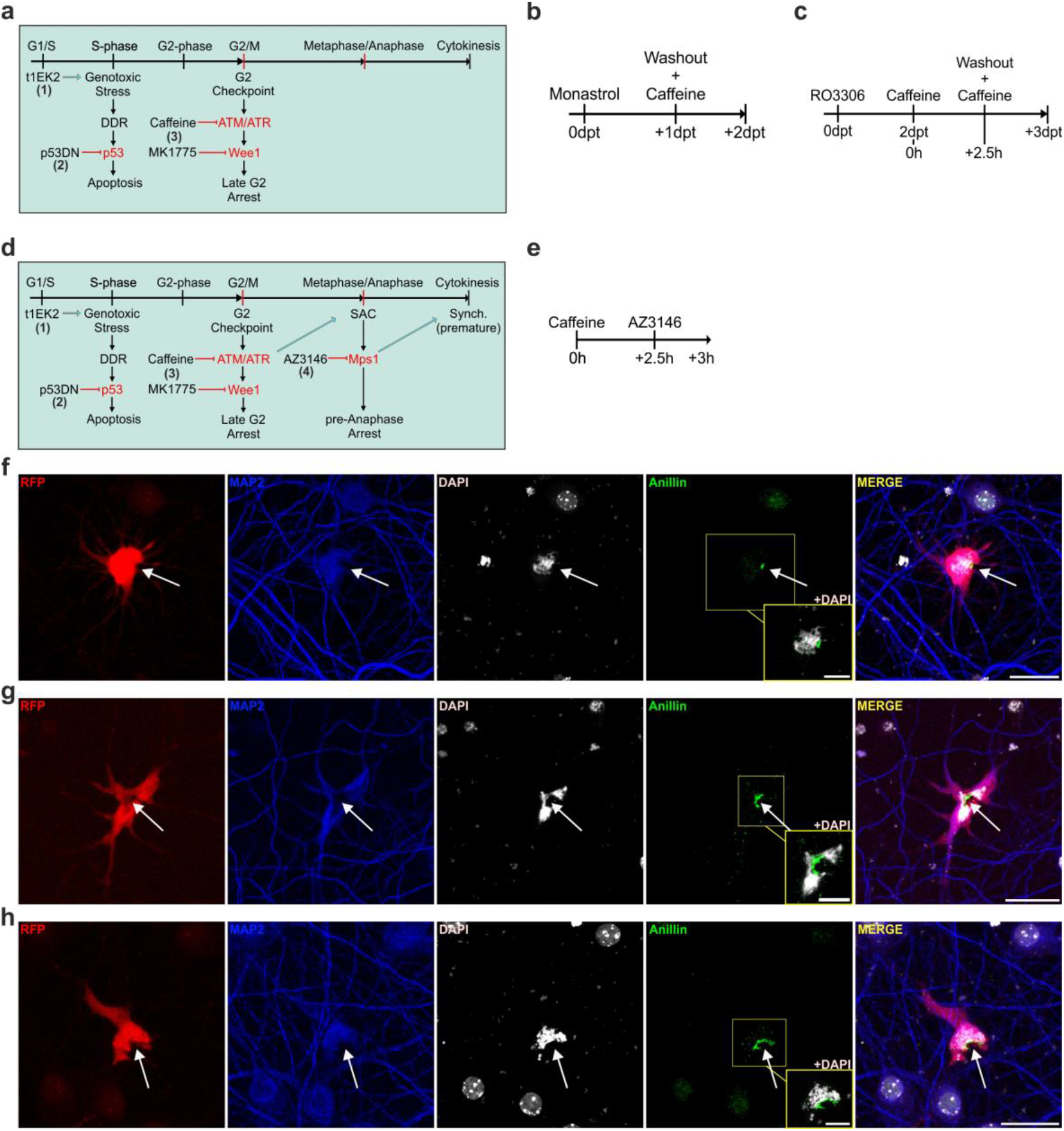
Differentiated neurons undergo cytokinesis. **a** Diagram extending on the regulation of M-phase and manipulations performed in this study shown in Fig. S1a. tEK2 induces G1/S transition (1)^18^. The loss of function of p53 affords viability and cell cycle progression (2). However, due to dysfunctional G1/S and S checkpoint regulation, p53-deficient cells rely heavily on G2 checkpoint arrest to amend DNA damage prior to M-phase entry^19^. Wee1 inhibition with MK1775 (3) can abrogate the G2 checkpoint^48^. Upstream of Wee1, the G2 checkpoint is dependent on ataxia telangiectasia-mutated (ATM) and ataxia telangiectasia and Rad3 related (ATR) to sustain G2 arrest^19^. Inhibition of ATM/ATR with caffeine (3) results in G2 checkpoint abrogation^50^. Insufficient decatenation of double stranded sister chromatid intertwines (dsSCI) can also delay metaphase entry (decatenation checkpoint) as well as anaphase onset^51^. Decatenation of dsSCI is topoisomerase 2 (TOP2) dependent. Thus, we expressed TOP2α^61^ to aid in the resolution of excess dsSCI that can potentially accumulate during an aberrant S-phase and lead to chromosome segregation errors^51^. **b** Protocol to induce cytokinesis at 1 dpt. The day of transfection, neurons were treated with the motor kinesin Eg5 inhibitor monastrol^65^ (100 μM) to prevent progression beyond prometaphase. Monastrol was used to prevent the progression of neurons that did not require abrogation of the G2 checkpoint to enter M-phase (Fig. 4a) because they are likely to have reduced levels of DNA damage (Fig. 4b, c, Video 1)^19^, which can facilitate M-phase completion^66^. At 1 dpt, monastrol was washed and caffeine (3 mM) was added to induce M-phase entry of neurons that were arrested by the G2 checkpoint. **c** Protocol to induce cytokinesis at 2dpt. The day of transfection, neurons were treated with the Cdk1 inhibitor R03306^67^ (9 μM) to prevent progression beyond late G2. RO3306 was preferred for longer cytokinesis induction experiments (2 dpt) because it cannot result in mitotic slippage^42^. Caffeine (3 mM) was added at 2 dpt to inhibit ATM/ATR^50^ before RO3306 washout to prime G2/M transition. Two and a half hours later RO3306 was washed and Caffeine (3 mM) was added again to sustain G2 checkpoint abrogation. **d** Diagram extending on Fig. S2a to include anaphase/cytokinesis synchronization (Synch.) by Spindle Assembly Checkpoint (SAC) abrogation for immunocytochemistry experiments. Once in M-phase, the SAC arrests progression in metaphase, preventing anaphase/cytokinesis onset until attachment between sister kinetochores and kinetochore microtubules is correct^20^. SAC activation prior to correct attachment requires active Monopolar spindle 1 (Mps1) targeting of Mitotic arrest deficient 2-like protein (Mad2) to kinetochores^22^. Mps1 inhibition with AZ3146 (4) can abrogate SAC response that requires kinetochore targeted Mad2^52^. Expression of TOP2α^61^ could also prevent additional non-kinetochore targeted Mad2-dependent anaphase delays owed to insufficient decatenation of dsSCI^68^. Noteworthy, insofar the SAC was not satisfied but abrogated, anaphase is inherently premature and thus the caveat is that synchronization can hinder anaphase/cytokinesis completion. **e** Protocol to induce cytokinesis for immunocytochemistry experiments. G2/M transition was induced with caffeine (3 mM). AZ3146 (10 μM) was added 2.5 h later to abrogate de SAC and induce anaphase/cytokinesis. Neurons were fixed for immunocytochemistry 30 min. later. **f-h** Confocal images of anillin immunolabelling of the onset of asymmetric cleavage furrow ingression (**f**) and progression (**g, h**). Cleavage furrow is pushing against the chromatin (**h**), likely reflecting that anaphase onset by SAC abrogation has been induced prematurely. Arrows identify RFP positive neurons. Bar: 25 μm.

## Video legends

**Video 1**. 3D reconstruction of interphase and prophase nuclei (DAPI, blue) in t1EK2/p53DN/RFP-expressing γ-H2AXpositive (not shown) neurons in Figure 4c.

**Video 2**. 3D reconstruction of RFP (red) and pH3 positive nucleus (green) of t1EK2/p53DN/RFP-expressing prometaphase-like neuron shown in Figure 4f.

**Video 3**. Left: Time-lapse video of t1EK2/p53DN/RFP/H2B-EGFP-expressing neuron remaining in interphase. Right: Time-lapse video of t1EK2/p53DN/RFP/H2B-EGFP-expressing neuron entering M-phase displaying dendritic alterations. At 1.5 dpt, neurons were treated with MG132 (10 μM) and 1 h later with MK1775 (900 nM) (Figure S1a, d). Time-lapse videos are at 1 frame per second (fps) taken at 20x magnification with intervals of 30 min.

**Video 4**. Time-lapse video of t1EK2/p53DN/TOP2α/RFP/H2B-EGFP-expressing neuron entering M-phase and attempting cytokinesis. The day of transfection, neurons were treated with monastrol (100 μM) to synchronize neurons at prometaphase. At 1 dpt, monastrol was washed out and neurons were treated with Caffeine (3 mM) to induce G2/M transition (Figure S2a, b). Time-lapse videos are at 1 fps taken at 20x magnification with intervals of 15 or 30 min.

**Video 5**. Time-lapse video of t1EK2/p53DN/TOP2α/RFP/H2B-EGFP-expressing neuron shown in Fig. 5a, b with intercellular bridge. The day of transfection, neurons were treated with monastrol (100 μM) to synchronize neurons at prometaphase. At 1 dpt, monastrol was washed out and neurons were treated with Caffeine (3 mM) to induce G2/M transition (Figure S2a, b). Time-lapse videos are at 1 fps taken at 20x magnification with intervals of 15 or 30 min.

**Video 6**. Time-lapse video of t1EK2/p53DN/TOP2α-RFP-expressing neuron shown in Fig. 5c, d completing division. The day of transfection, neurons were treated with RO3306 (9 μM) to synchronize neurons at late G2. At 2 dpt, neurons were treated with caffeine (3 mM) to inhibit ATR/ATM. RO3306 was washed out 3.5 h later to release neurons from late G2 arrest and caffeine (3 mM) was added again to induce G2/M transition (Figure S2a, c). Time-lapse images are taken at 20x magnification at 1 fps with each frame taken at intervals of 30 min.

**Video 7**. 3D reconstruction of RFP (red), MAP2 (grey) anillin (green) and DAPI (blue) of t1EK2/p53DN/TOP2α/RFP-expressing neuron shown in Figure 5f (see protocol in Figure S2d, e).

**Video 8**. 3D reconstruction of RFP (red), pH3 (green) and DAPI (blue) of t1EK2/p53DN/TOP2α/RFP-expressing neuron shown in Fig. 5i (see protocol in Figure S2d, e).

